# Epigenetic clocks reveal a rejuvenation event during embryogenesis followed by aging

**DOI:** 10.1101/2021.03.11.435028

**Authors:** Csaba Kerepesi, Bohan Zhang, Sang-Goo Lee, Alexandre Trapp, Vadim N. Gladyshev

## Abstract

The notion that germline cells do not age goes back to the 19^th^ century ideas of August Weismann. However, being in a metabolically active state, they accumulate damage and other age-related changes over time, i.e., they age. For new life to begin in the same young state, they must be rejuvenated in the offspring. Here, we developed a new multi-tissue epigenetic clock and applied it, together with other aging clocks, to track changes in biological age during mouse and human prenatal development. This analysis revealed a significant decrease in biological age, i.e. rejuvenation, during early stages of embryogenesis, followed by an increase in later stages. We further found that pluripotent stem cells do not age even after extensive passaging and that the examined epigenetic age dynamics is conserved across species. Overall, this study uncovers a natural rejuvenation event during embryogenesis and suggests that the minimal biological age (the ground zero) marks the beginning of organismal aging.

---

Aging is characterized by a progressive accumulation of damage, leading to loss of physiological integrity, impaired function and increased vulnerability to death (*1*). While the aging process affects the entire organism, it is often discussed that the germline does not age, because this lineage is immortal in the sense that the germline has reproduced indefinitely since the beginning of life (*2–4*). This notion dates to the 19th century when August Weismann proposed the separation of ageless germline and aging soma. However, being in the metabolically active state for two decades or more before its contribution to the offspring, human germline accumulates molecular damage, such as modified long-lived proteins, epimutations, metabolic by-products, and other age-related deleterious changes (*5*, *6*). It was shown that sperm cells exhibit a distinct pattern of age-associated changes (*7–9*). Accordingly, it was recently proposed that germline cells may age and be rejuvenated in the offspring after conception (*10,11*). If this is the case, there must be a point (or period) of the lowest biological age (here, referred to as the ground zero) during the initial phases of embryogenesis (Fig. 1A). Here, we carried out a quantitative, data-driven test of this idea.

**Fig. 1.**
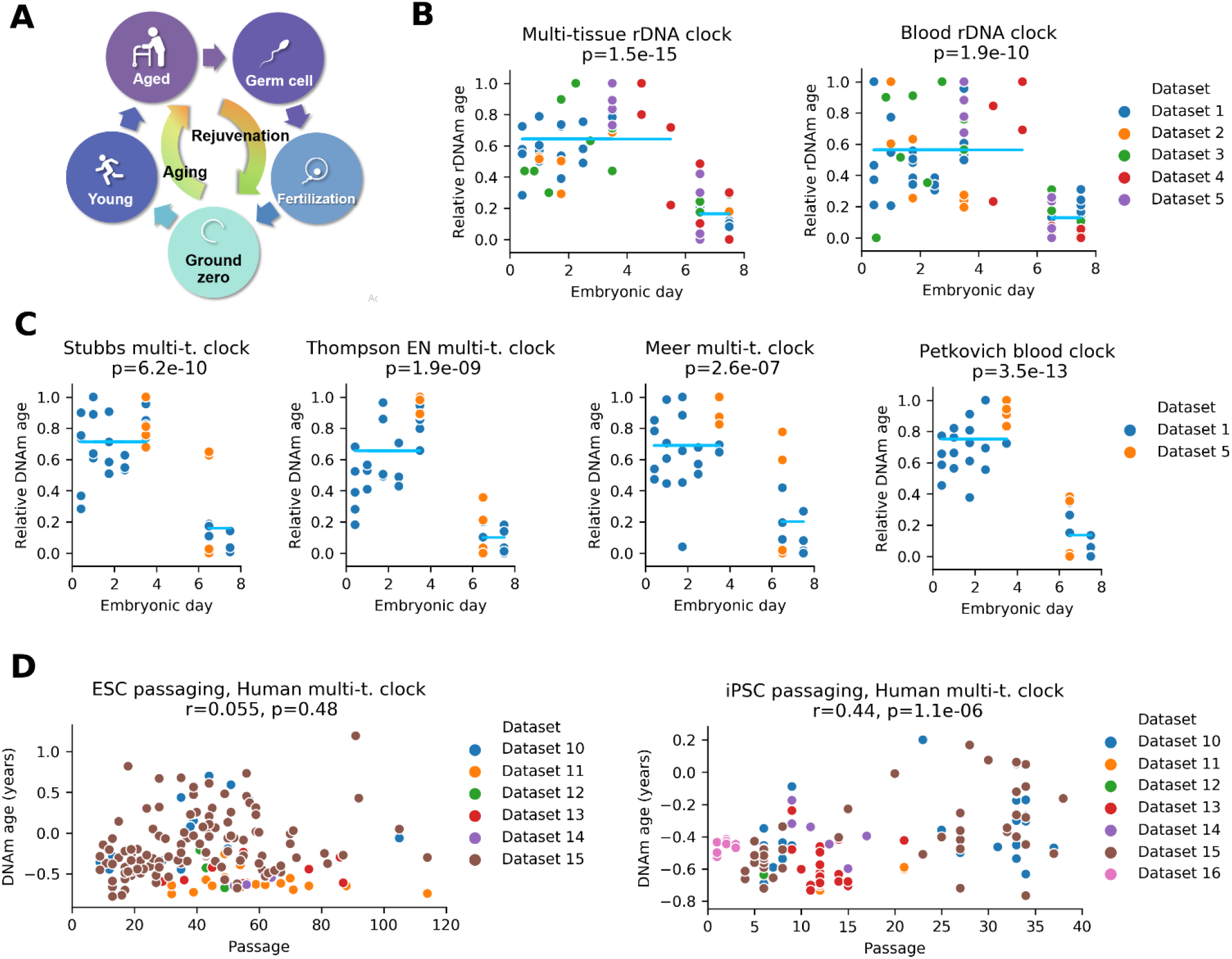
A rejuvenation event during early embryogenesis revealed by aging clocks. **(A)** Overview of the model, which posits that germline cells age during development and adulthood and are rejuvenated in the offspring after conception. The model also suggests that there is a time point corresponding to the lowest biological age (ground zero). **(B)** Multi-tissue and blood rDNA clocks applied to five datasets spanning the first 8 days of mouse embryogenesis (Table 1, datasets 1-5). We rescaled epigenetic age of each dataset to the interval [0,1] for comparison (‘relative rDNA age’). 0 represents the lowest epigenetic age and 1 represent the highest epigenetic age of each dataset. Blue lines indicate the mean of each group; p-values of two-sided t-test comparing the means of the two groups (before and after E6) are displayed. **(C)** Application of four genome-wide epigenetic aging clocks to two available mouse RRBS datasets. **(D)** Epigenetic age of human ESCs and iPSCs as a function of passage number. Horvath human multi-tissue clocks were applied.

Due to recent advances in technology, machine learning is flourishing and has led to breakthroughs in many areas of science by discovering multivariate relationships (*12*). Aging and developmental biology areas also exploited the potential of machine learning by developing algorithms (“aging clocks”) that can estimate chronological age or biological age (i.e. the age based on molecular markers) of an organism from a given data (*13, 14*). As epigenomic changes, which result in dysregulation of transcriptional and chromatin networks, are crucial components of aging (*15*), epigenetic clocks, based on methylation levels of certain CpG sites, emerged as a promising molecular estimator of biological age (*16*, *17*). These clocks were shown to quantitatively measure numerous aspects of human aging (*17*–*22*). For example, epigenetic age acceleration was associated with age-related conditions, such as all-cause mortality (*23*, *24*), cognitive performance (*25*), frailty (*26*), Parkinson’s disease (*27*), Down syndrome (*28*), and Werner syndrome (*29*). Epigenetic aging clocks were also developed for mice and could be used to evaluate longevity interventions, such as calorie restriction and growth hormone receptor knockout (*30–36*). Most importantly, the clocks developed based on aging patterns of mostly adult tissues report the effects of cell rejuvenation upon complete- or partial reprogramming of adult fibroblasts into induced pluripotent stem cells (iPSCs) as demonstrated in both human and mouse (*33*, *35*, *37–39*). Even though iPSCs correspond to an embryonic state and the transition to these cells involves major molecular changes, including changes in the epigenome, this rejuvenation event can be assessed by epigenetic clocks. Most recently, universal mammalian clocks have been developed based on conserved cytosines, whose methylation levels change with age across mammalian species (*40*).

Considering that epigenetic clocks track the aging process, they may be applied to early development to characterize biological age dynamics during that period of life. Recent studies showed that clocks may be successfully applied to human fetal development using brain, retina, and cord blood samples (*41*–*43*). However, epigenetic age dynamics during entire prenatal development for the entire organism remained unexplored. Here, we developed a new multi-tissue epigenetic clock using machine learning and applied it, together with other existing aging clocks, to assess prenatal development in mammals from the perspective of aging. This approach uncovered a rejuvenation period during early embryogenesis and the timing of the beginning of aging in mammals.

## A rejuvenation event during early embryogenesis

To assess epigenetic age dynamics during embryogenesis, we collected available human and mouse DNA methylation datasets (Table 1) and subjected them to various epigenetic aging clocks (Table 2). We also developed a multi-tissue ribosomal DNA methylation clock (rDNAm) (fig. S1, S2). The rDNA is characterized by a large number of age-associated CpG sites that exhibit high sequence coverage due to the multiplicity of rDNA in the genome (*35*, *44*). The new clock is capable of predicting the epigenetic age of RRBS (reduced representation bisulfite sequencing), WGBS (whole-genome bisulfite sequencing) and even pseudo-bulk single cell sequencing samples in various tissues. All clocks we employed showed high accuracy (r >= 0.8) in age prediction of test samples and were sensitive to age-related conditions and longevity interventions (Table 2).

**Table 1.** Embryonic DNA methylation datasets used in this study.

| Name | Reference | Accession | Platform | Species | Samples (number of biological replicates) |
| --- | --- | --- | --- | --- | --- |
| Dataset 1 | (56) | GSE34864 | RRBS | mouse | Zygote (5), 2-cell (4), 4-cell (5), 8-cell (3), ICM (5)<br>E6.5 (4), E7.5 (5) |
| Dataset 2 | (49) | GSE56697 | WGBS | mouse | 2-cell (2), 4-cell (2), E3.5 (3), E6.5 (2), E7.5 (2),<br>E13.5 (2) |
| Dataset 3 | (57) | GSE98151 | WGBS | mouse | Zygote (1), early 2 cell (1), late 2 cell (1), 4 cell (1),<br>8 cell (1), morula (1), ICM (1), trophoblast (1),<br>E6.5 epiblast (1), E6.5 ext ect (1), E7.5 epiblast (1),<br>E7.5 ext ect (1) |
| Dataset 4 | (55) | GSE121690 | scNMT-seq | mouse | 758 single cells from E4.5, E5.5, E6.5, E7.5 |
| Dataset 5 | (61) | GSE51239 | RRBS | mouse | ICM (2), trophectoderm (2), E6.5 epiblast (2), E6.5<br>ext end (2), ESC P0-P5 (17) |
| Dataset 6 | (59) | ENCSR486XIX | WGBS | mouse | Various tissues from E10.5 to birth (139) |
| Dataset 7 | (62) | GSE56515 | 450K array | human | Various tissues from GW 9 to GW 22 (34) |
| Dataset 8 | (63) | GSE31848 | 450K array | human | Various tissues from GW 14 to GW 20 (37) |
| Dataset 9 | (64) | GSE69502 | 450K array | human | Various tissues from GW 14.5 to 23. (49) |
| Dataset 10 | (63) | GSE31848 | 450K array | human | ESC P9 - P105 (19), iPSC P5 - P37 (29) |
| Dataset 11 | (65) | GSE34869 | 450K array | human | ESC P32 - P114 (19), iPSC P12 - P21 (5) |
| Dataset 12 | (66) | GSE40909 | 450K array | human | ESC P41 - P49 (3), iPSC P6 (2) |
| Dataset 13 | (67) | GSE44424 | 27K array | human | ESC P29 - P87 (8), iPSC P9-P21 (21) |
| Dataset 14 | (68) | GSE51747 | 27K array | human | ESC P52 - P64 (3), iPSC P9-P17 (6) |
| Dataset 15 | (63) | GSE30653 | 27K array | human | ESC P9 – P114 (116), iPSC P4-P69 (46) |
| Dataset 16 | (69) | GSE54848 | 450K array | human | iPSC P1-P3 (9) |

E, gestational day; ESC, embryonic stem cell; P, passage; ext ect, extraembryonic ectoderm; ext end, extraembryonic endoderm; GW, gestational weeks; iPSC, induced pluripotent stem cell; RRBS, Reduced representation bisulfite sequencing; WGBS, whole genome bisulfite sequencing

**Table 2.**
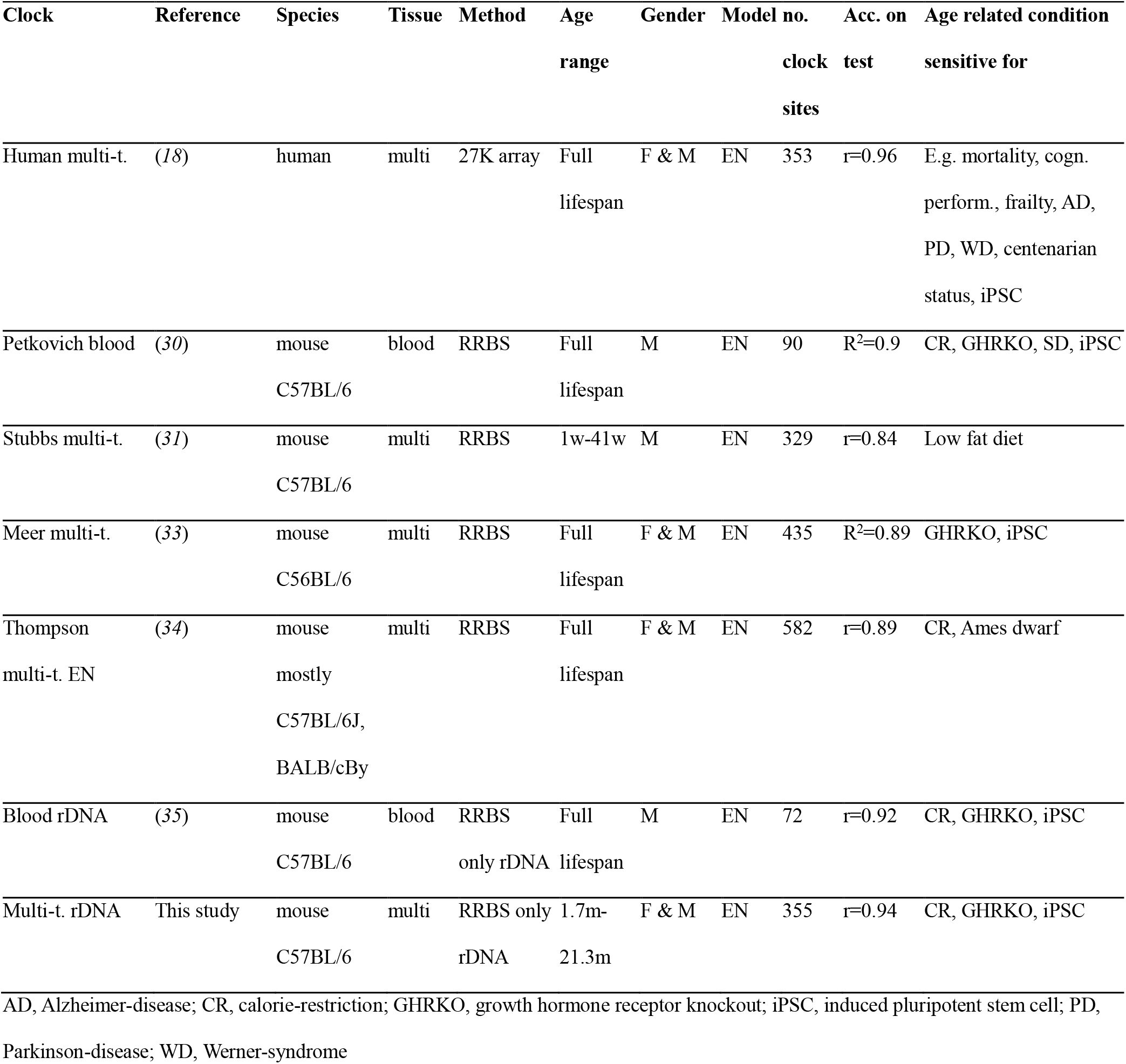
Epigenetic aging clocks used in this study.

First, we examined the behavior of rDNAm clocks when applied to five independent mouse early embryonic datasets. We found that the mean epigenetic age of E6.5/E7.5 embryos was consistently lower than in earlier stages of embryogenesis (Fig. 1B, fig. S3AB, fig. S4AB). We also applied four genomewide RRBS-based epigenetic aging clocks to RRBS datasets (datasets 1 and 2). This again revealed that the epigenetic age of E6.5/E7.5 embryos is lower than during the period from zygote to blastocyst (Fig. 1C, fig. S3CD, fig. S4CD). Thus, epigenetic age decreases during early embryogenesis, and therefore embryonic cells not only do not age during this period but at some point get rejuvenated.

Previously, a near-zero epigenetic age of human embryonic stem cells and iPSCs was demonstrated by the Horvath multi-tissue clock even after extensive passaging (*18*). We analyzed iPSCs and ESCs based on several currently available datasets to further assess whether these cells, which correspond to early embryogenesis, age (Fig. 1D). The epigenetic age of cells was very low (mostly below zero) even after more than 100 passages. Even under artificial culture conditions, at the level of oxygen above physiological, and with the number of passages well beyond physiological (which may lead to the accumulation of deleterious mutations), either no or very little increase in epigenetic age was observed. These findings support the notion that cells corresponding to the early stages of embryogenesis essentially do not age.

## Organismal aging begins during mid-embryonic development in mouse and human

We quantified the epigenetic age by applying rDNA clocks to the only available mouse dataset that contains both early- and late embryonic samples (Fig. 2A). The epigenetic age at E6.5 and E7.5 was significantly lower than at E13.5 (primordial germline cells that are the direct progenitors of sperm and oocytes).

**Fig. 2.**
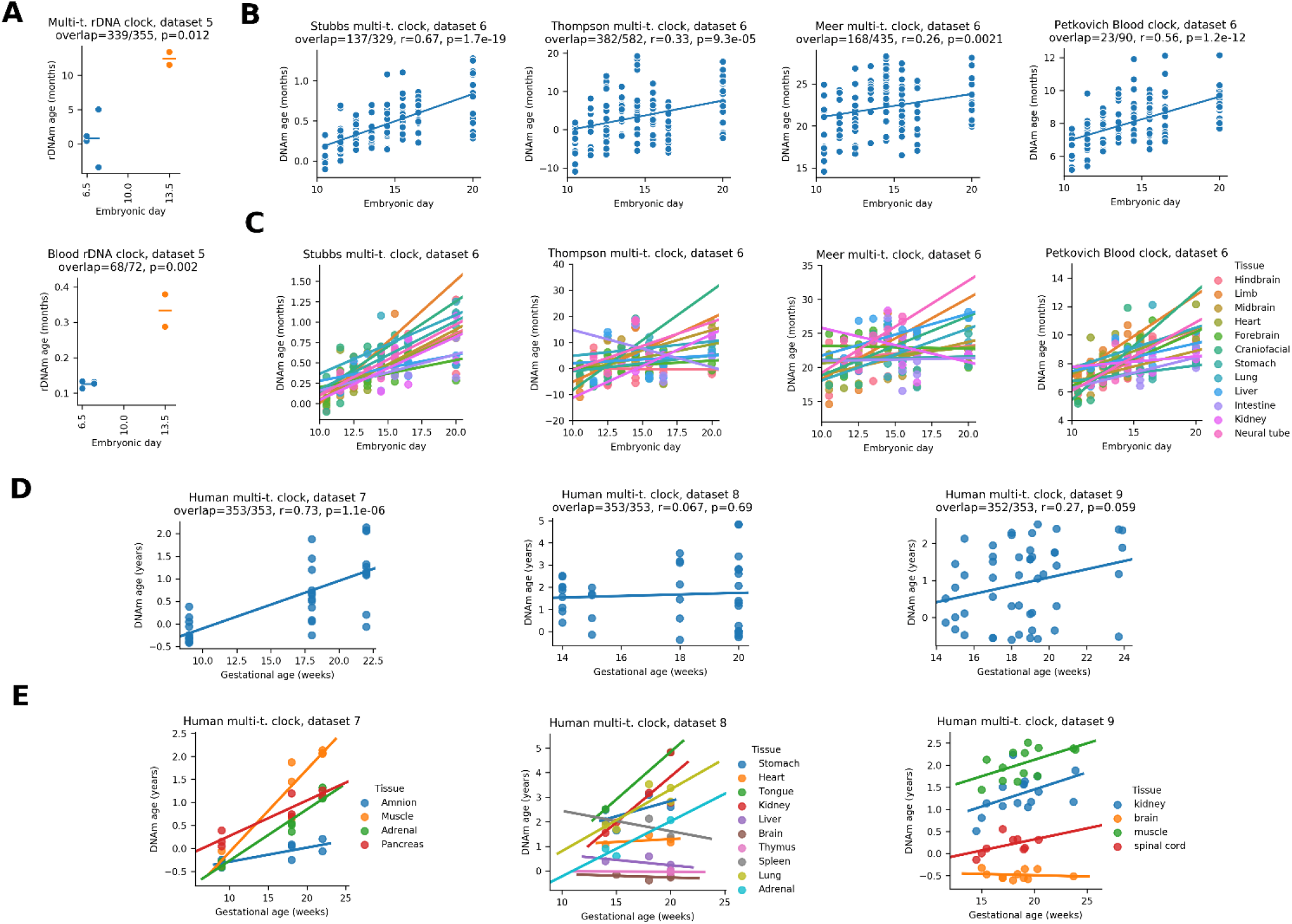
Organismal aging begins in mid-embryonic development in mouse and human. **(A)** Epigenetic age (multi-tissue and blood rDNA clocks) analysis of the dataset that contains both early and late mouse embryo samples (E13.5 samples are based on primordial germline cells). **(B)** Application of genome-wide epigenetic clocks to later stages of mouse embryogenesis (r, Pearson correlation coefficient; p, p-value of the correlation). **(C)** The same data as above, but separated by tissue. An increasing trend is observed for almost all tissues, with few non-significant exceptions. **(D)** Epigenetic age dynamics of four independent prenatal human 450k methylation array datasets based on the Horvath human multi-tissue clock. **(E)** The same data as above, but separated by tissue (5 significant increases, 9 non-significant increases, 4 non-significant decreases, 0 significant decreases).

We also assessed the epigenetic age of mouse embryos across 9 time points from embryonic day 10.5 to birth (dataset 6) by genome-wide methylation clocks. A consistent increase in epigenetic age was observed during this period both when considering all data (Fig. 2B) and when separated by tissue (Fig. 2C). In addition, we analyzed human prenatal datasets by applying the Horvath multi-tissue DNAm clock to four independent human 450K methylation array datasets (datasets 7-9). An increase (or in some cases no change) in epigenetic age was observed both when considering all data (Fig. 2D) and tissue-by-tissue (Fig. 2E). Thus, at a certain point during embryonic development in mouse and human the biological age begins to increase in most or all tissues. Considering that epigenetic clocks track the aging process, the data suggest that by then organisms already age.

## Epigenetic age of mouse ESCs during early passaging

We assessed the epigenetic age of mouse embryonic stem cells after outgrowth (passage 0) and early passaging (passage 5) under three different culture conditions (Fig. 3AB). In the absence of two inhibitors (PD0325901 that causes blockade of differentiation and CHIR99021 that supports self-renewal (*45*)), we observed a lower epigenetic age after outgrowth compared to the condition when the two inhibitors were included. The data suggest that ESCs under incomplete self-renewal culture conditions may continue their development (without self-renewal) and rejuvenate, similar to what we observed *in vivo* (Fig. 1).

**Fig. 3.**
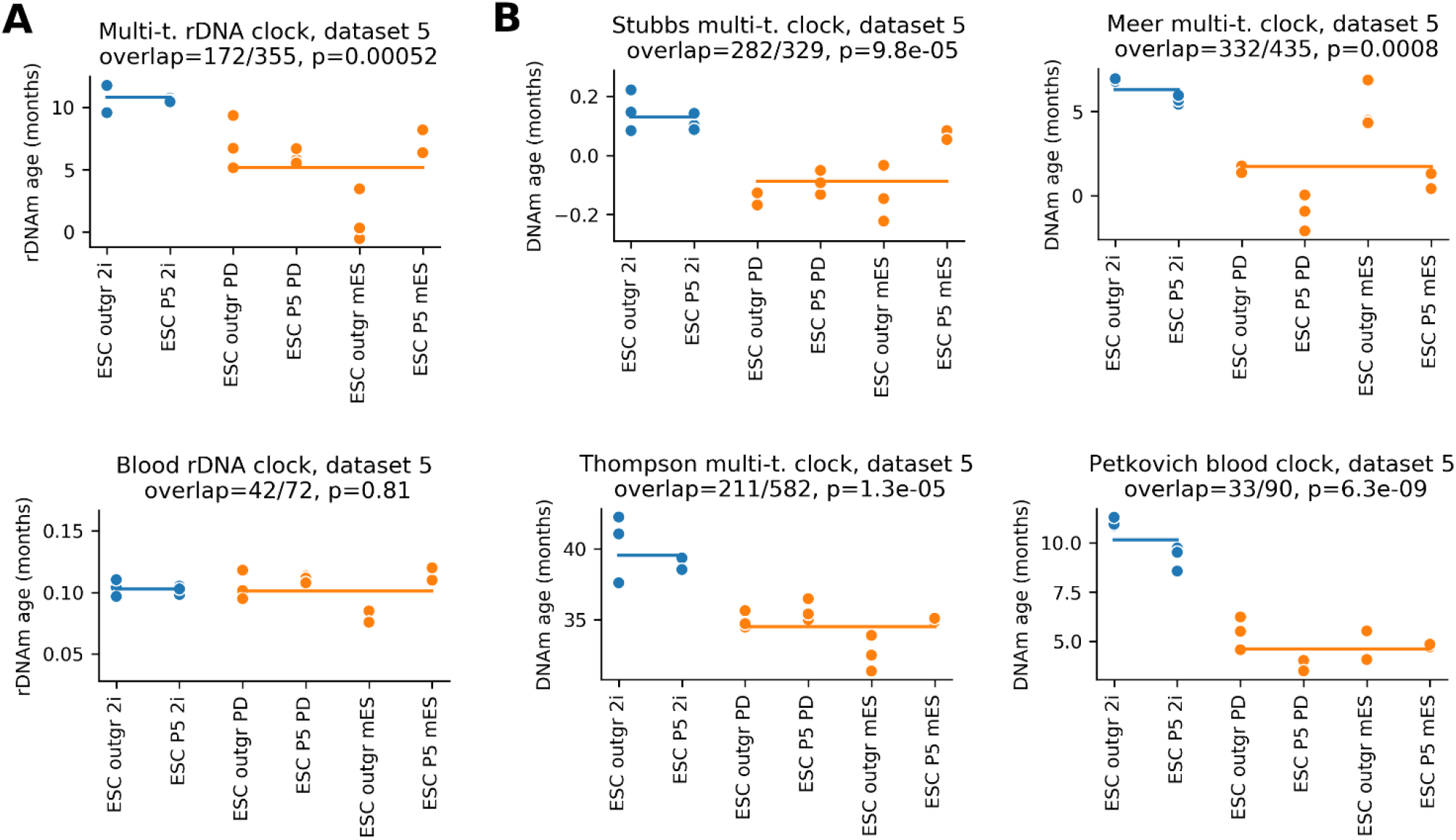
Epigenetic age of mouse ESCs during early passaging under different culture conditions. **(A)** Epigenetic age (by rDNA clocks) of mouse embryonic stem cells after outgrowth (passage 0) and passage 5 under three different culture conditions (2i, both self-renewal supporting inhibitors used; only one inhibitor; mES, no inhibitor). **(B)** Application of genome-wide mouse epigenetic clocks to the same data.

## Localization of the epigenetic age minimum (ground zero) during mouse embryonic development

We concatenated the results for early and later stages of embryogenesis by applying genome-wide mouse epigenetic clocks (Fig. 4). The variable number of overlapped clock sites across all stages caused a batch effect that resulted in a shift of the actual predicted age between early and late stages. However, the epigenetic age dynamics showed a clear U-shaped pattern in every case, with the minimum at E7.5 in three cases and E10.5 in one case. The exact localization of the minimum was not possible with the data currently available, and it may lie in the range from E4.5 to E10.5. The data suggest that organismal aging begins at that period after the rejuvenation event.

**Fig. 4.**
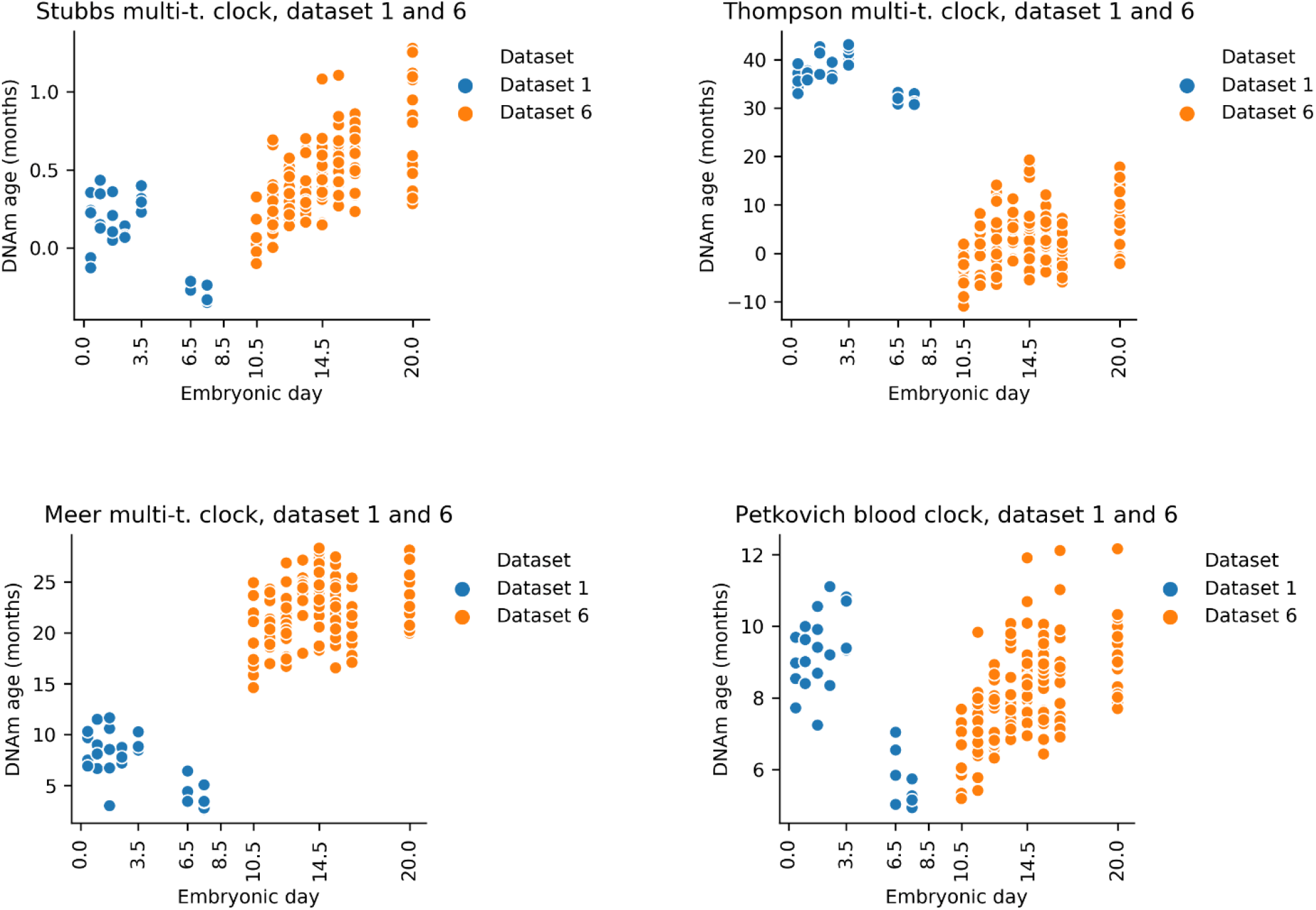
Localization of the epigenetic age minimum (ground zero) during mouse embryonic development. We concatenated results for entire embryogenesis by using the genome-wide mouse epigenetic clocks indicated. In all cases, the epigenetic age minimum is observed during embryogenesis.

## Concluding comments

Back in the 19^th^ century, August Weismann proposed the idea of heritable non-aging germline and disposable aging soma. Yet, the germline shows molecular changes characteristic of aging (*7–9*). Our study suggests that the germline ages but is rejuvenated in the offspring at some point during early embryogenesis. This rejuvenation occurs during early post-implantation stages when the offspring reaches its minimal biological age. We propose that this minimum, the ground zero, marks the beginning of aging of an organism. The offspring proceeds naturally to ground zero from the zygote stage, but somatic cells may also be forced to this young stage, e.g. by reprogramming with Yamanaka factors (or by other methods), generating iPSCs. In vivo amelioration of age-associated hallmarks was already demonstrated by partial reprogramming (*46*). Most recently, partial reprogramming restored vision in mice by resetting youthful epigenetic information (*47*). Thus, both soma and germline may age and be rejuvenated.

Early embryogenesis, where we observed a rejuvenation period, is also accompanied by other molecular changes in preparation for organismal life, such as a gradual extension of telomeres (*48*), waves of global demethylation and methylation (*49*), transition from the use of maternal gene products to those of the embryo, inactivation of chromosome X, and development of monoallelic gene expression (*50*). Rejuvenation should also involve a decrease in molecular damage and other deleterious age-related changes that accumulate in the parental germline (*51, 52*). The data indicate that ground zero lies between E4.5 and E10.5 in mice, and the current estimates suggest that it is close to E6.5/E7.5. This period approximately corresponds to gastrulation, where three germ layers are formed. However, further studies are needed to precisely localize ground zero in humans and mice.

The beginning of aging is a subject to debate. It is often discussed that aging begins after completion of development, at the onset of reproduction, and at the time when strength of natural selection begins to decrease. However, our recent analysis of deleterious age-related changes revealed that aging begins early in life, even before birth (*53*). Our current work now pinpoints the beginning of aging to ground zero.

CpG sites associated with aging and lifespan may be both hypomethylated and hypermethylated upon aging (*54*). An attractive possibility is that rejuvenation may be supported by remethylation rather than by demethylation. Indeed, global remethylation was reported between E3.5 and E7.5 (*49*, *55*–*57*), which is the same period where we observe rejuvenation. If global remethylation is indeed associated with epigenetic age decrease, ground zero and global methylation maximum should correspond to the same developmental stage. This would make sense from the perspective that, in order to remove “epigenetic damage”, the genome should be first partially demethylated and then remethylated again.

We also found that cells corresponding to early embryogenesis, i.e. ESCs and iPSCs, do not age when cultured and passaged. However, early passaging seems to result in epigenetic age reduction. Consistent with this age reduction, it was found that initial passaging induces telomere extension, and that mice generated from these rejuvenated cells live longer and are better protected from age-related diseases than the mice from the same cells that were not passaged (*58*). This suggests an exciting possibility that the natural rejuvenation event we uncover in this work may be targeted, such that organisms may begin aging at a lower biological age and therefore may achieve longer lifespan and extended healthspan. This may also be useful during *in vitro* fertilization, wherein embryos with a lower biological age may be prioritized.

Global cytosine methylation (average methylation level of CpG sites representing the whole genome) changes in waves during mammalian embryogenesis: an initial decrease from zygote to E3.5 is followed by an increase to E6.5/E7.5 (*49*), and unique tissue-specific dynamics during/after organogenesis (*59*). However, global cytosine methylation shows very little change after birth (*60*), and therefore is not highly predictive of biological age or its reduction. In contrast, the age predicted by epigenetic aging clocks (usually based on several hundred CpG sites) shows strong correlation with age (r >= 0.8), indicating it can be used to predict biological aging and rejuvenation (Table 2).

Overall, this work identifies a natural rejuvenation event during early life and suggests that organismal aging begins during embryogenesis, approximately at the time of gastrulation. These findings provide opportunities for understanding what this early rejuvenation process entails, whether it is similar to the Yamanaka reprogramming, and whether it may be induced in somatic cells in order to rejuvenate them.

## Acknowledgements

Authors thank Sun Hee Yim, Anastasia Shindyapina, Margarita Meer, Marco Mariotti, Maxim Gerashchenko for discussion. Supported by NIH grants to V.N.G.

## Author contributions

V.N.G. conceived and supervised the study. C.K. acquired data, performed data analysis and developed the multi-tissue rDNA clock. B.Z. created the schematic figure and helped with data analysis and discussions. All authors interpreted the results. V.N.G. and C.K. wrote the manuscript. B.Z., S.L., and A.T edited the manuscript.

## Data Availability and Code Availability

Methylation levels of rDNA CpG sites will be available for all mouse datasets upon publication along with the codes of epigenetic clock workflow.

## Supplementary Information

### Acquisition and processing of sequencing datasets

Raw sequences were downloaded and extracted using SRA Toolkit. Reads were trimmed and quality filtered by TrimGalore! v0.6.4 using the --rrbs option for RRBS. Methylation levels were extracted using Bismark v0.22.2(*70*) in two separate processes: (i) for rDNA clocks, reads were mapped to the mouse rDNA sequence (BK000964.3), (ii) for genomic clocks, reads were mapped to the complete mouse genome sequence (mm10/GRCm38.p6). Methylated and unmethylated cytosines of technical repeats were summed for each position. In the case of dataset 6, we directly used the available BED files (URLs were extracted from GEO files).

### CpG site filtering and imputation of RRBS/WGBS datasets

Our filtering strategy was optimized for three competing goals: (i) highest possible read coverage; (ii) highest possible clock site overlap; and (iii) best (unbiased) sample comparison. We used only the CpG sites that are covered by at least 5 reads (or 50 in the two highest covered cases when rDNA clocks were applied on datasets 2 and 3) in at least 90% of the samples of a given dataset. In the case of dataset 6, we just used the CpG sites that are covered by at least 5 reads without any other restriction (to avoid too many missing values). We omitted the two lowest quality samples (both were oocytes) of dataset 1 for genomic analysis. Missing values of clock sites were imputed by the average methylation levels of the covered clock sites for all samples. Our imputation strategy resulted in a single number for each clock and dataset and this allowed a less biased comparison of the samples compared to a method that imputes different values for each sample. We used a clock-site-wise average instead of all-site-wise average because we supposed that methylation levels of clock CpG sites are closer to each other than to other ones.

### Application of mouse epigenetic clocks to RRBS/WGBS datasets

In the application of previously developed clocks, we followed the descriptions of authors. All available mouse clocks that we applied were based on linear regression (Table 2). We used the intercept and weights provided by the authors. Transformation was also applied in the case of Petkovich blood clock (*30*), Stubbs multi-tissue clock (*31*) and blood rDNA clock (*35*). In the case of the Stubbs multi-tissue clock, an initial normalization was applied by the published training set. Applying RRBS-based clocks with sites scattered across the genome is often challenging, because of a varying number of covered clock sites, leading to missing values. Therefore, we required the clock site coverage to be at least 25%. RRBS-based clock sites had no sufficient coverage when we applied them to datasets 2, 3 and 4 (all of them are WGBS datasets).

### Single cell clock workflow

Methylation levels were obtained by the rDNA mapping workflow described above. In the end, we obtained methylation data for each of the 758 cells separately. We used only the CpG sites that were covered by at least 5 reads and showed consistent methylation levels (higher than 0.8 or lower than 0.2). Then, we randomly assigned cells to two groups, for each embryo stage (8 cell groups total). We calculated the average methylation status for each CpG site of each of the 8 groups (previously, methylation status was re-assigned to 0 or 1). Finally, we applied rDNA clocks to the 8 groups as it would be 8 samples from bulk sequencing.

### Application of human DNAm clock to 450K methylation array datasets

Datasets were downloaded from GEO database. We re-indexed data to the 27K array and used the R script of the Horvath DNAm calculator as described in the tutorial (https://horvath.genetics.ucla.edu/html/dnamage/TUTORIAL1.pdf). In the case of PSC datasets, we only analyzed ESCs derived from normal fertilized blastocysts and iPSCs derived from normal cells (i.e. immortalized cells were excluded).

### Development of a mouse multi-tissue rDNA clock

A multi-tissue mouse RRBS dataset (GSE120132) was used for training and testing the clock by crossvalidation. An additional blood mouse RRBS dataset (GSE80672) was used for testing samples of an independent study. Raw sequences were downloaded from the SRA database. Reads were trimmed and quality filtered by TrimGalore! v0.6.4, and methylation levels were extracted using Bismark v0.22.2. by mapping to the mouse rDNA sequence (BK000964.3). Fastq files of technical repeats were concatenated. We used only the CpG sites (and only positive-strand cytosines) that are covered in at least 50 reads in all of the samples of both datasets. We used only the C57BL/6J of the GSE120132 dataset and omitted muscle samples because we observed low age correlation of the CpG sites of muscle rDNA (fig. S5). After filtering, 166 samples remained for training and testing (fig. S2A). Then, 5-fold cross-validations were performed on all of the 166 samples for different lambda parameters of ElasticNet (Scikit-learn v0.23.2) to find the optimal lambda parameter (fig. S1B). We re-trained a final model (the actual clock) on the same dataset with the optimal parameter (lambda=0.0001) on the random 80% of all samples and tested on the remaining samples (fig. S1C, table S1). Our final model showed a similar performance to the average performance of the cross-validation. However, testing a model on the dataset of the cross-validation may lead to inflated performance. To resolve this issue, we also examined our model in an independent test. That is, we tested the multi-tissue clock on 153 of normal (control-fed, wild type) C57BL/6J blood samples (fig. S1C). We also applied it for age-related conditions (fig. S2).

---

**Figure S1.**
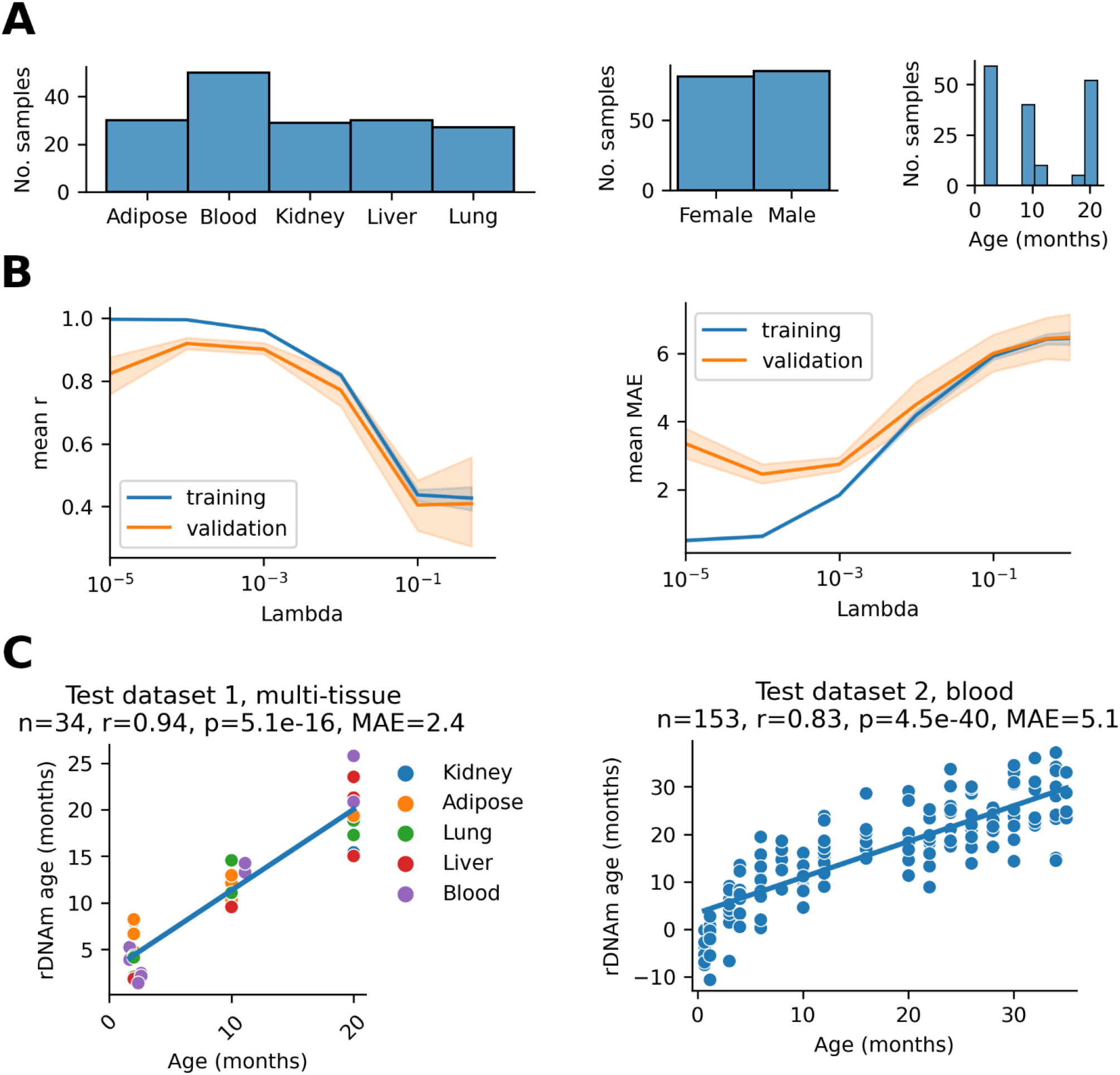
Development of a mouse multi-tissue ribosomal DNA methylation (rDNA) clock. **(A)** Sample distribution used for training and testing. **(B)** Finding the optimal lambda parameter (0.0001) of ElasticNet using 5-fold cross-validation. **(C)** Performance of the multi-tissue rDNA clock on a multi-tissue test set and a blood test set, both with normal/control conditions.

**Figure S2.**
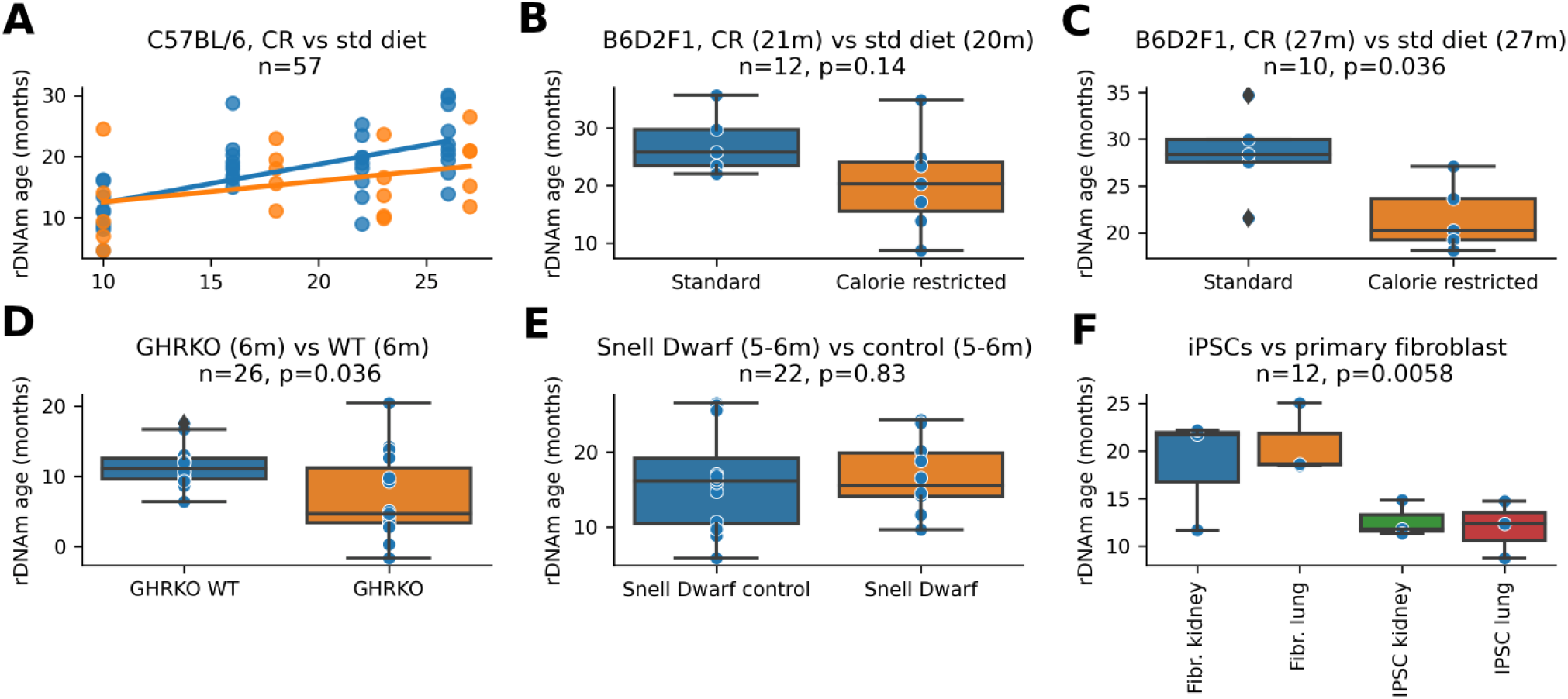
Application of the multi-tissue rDNA clock to various models of aging and to longevity interventions. **(A)** Calorie restricted (orange) vs standard diet fed (blue) C57BL/6 mice. **(B)** Calorie restricted 21 months old vs standard diet fed 20 months old B6D2F1 mice. **(C)** Calorie restricted 27 months old vs standard diet fed 27 months old B6D2F1 mice. **(D)** Growth hormone receptor knockout mice (6 months old) vs wild type mice (6 months old). **(E)** Snell dwarf 5-6 months old vs control mice. **(F)** Mouse lung and kidney fibroblasts vs fibroblast-derived induced pluripotent stem cells (iPSCs).

**Figure S3.**
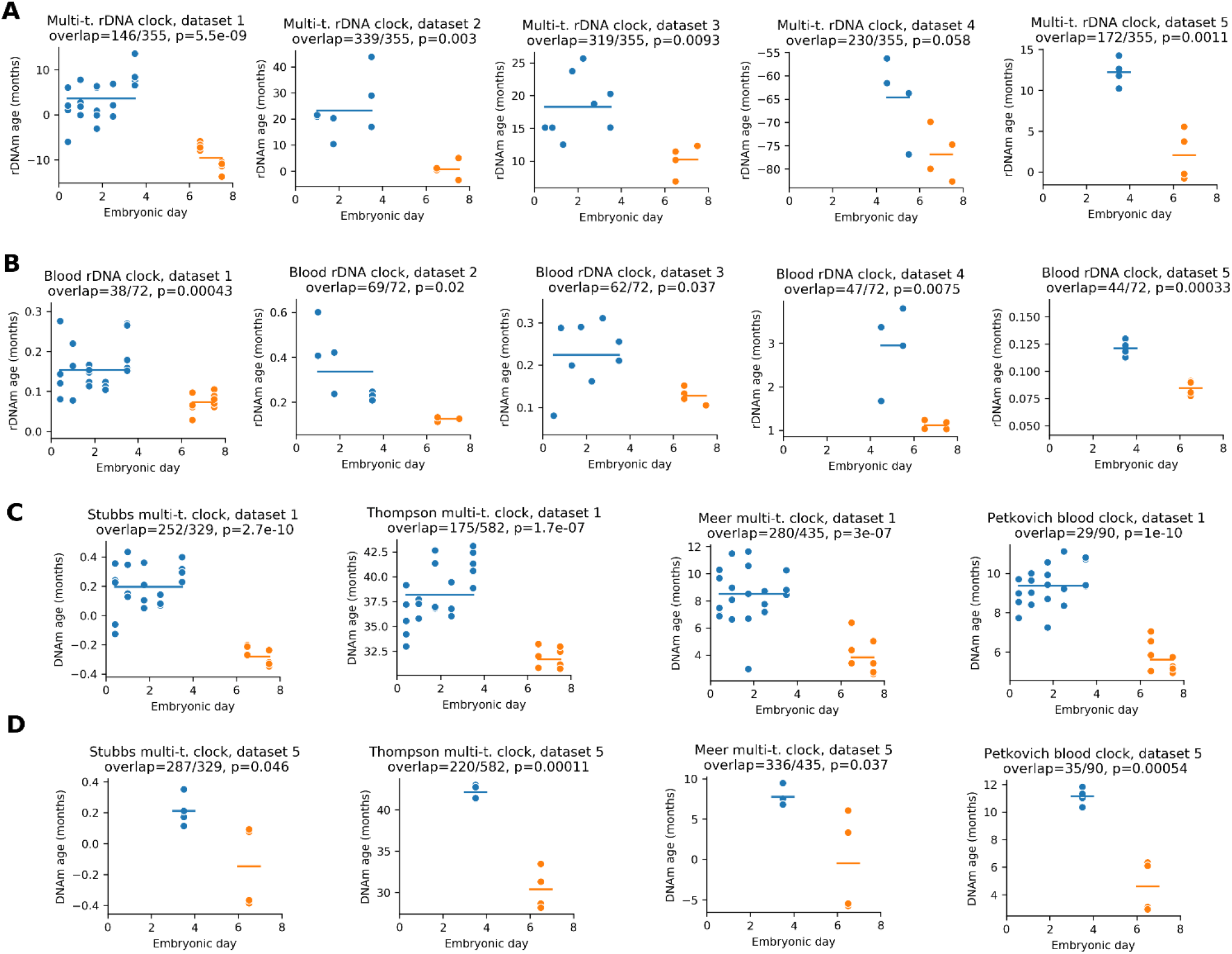
Aging clocks reveal a rejuvenation event during early embryogenesis. **(A)** Multi-tissue and blood rDNA clocks applied to five datasets spanning the first 8 days of mouse embryogenesis (Table 1, datasets 1-5). Predicted epigenetic age is displayed (no rescaling is applied). For each plot, we indicate the number of clock CpG sites that were covered in the application dataset (‘overlap’). Blue lines indicate the mean of each groups; p-value of two-sided t-test compares the mean of the two groups. **(B)** Blood rDNA methylation clock applied to the same five datasets. **(C)** Application of four genome-wide epigenetic aging clocks to dataset 1. **(D)** Application of four genome-wide epigenetic aging clocks to dataset 5.

**Figure S4.**
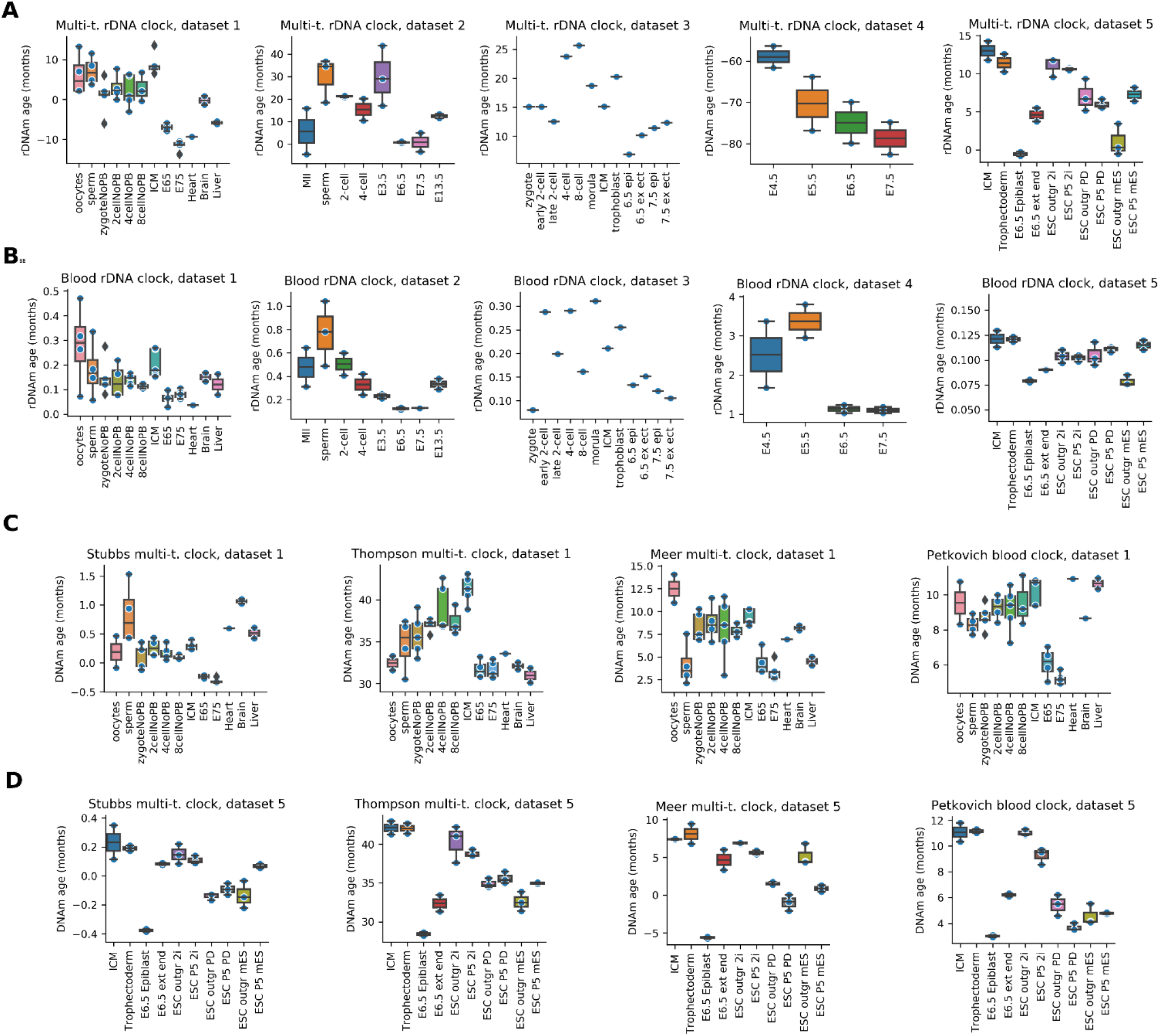
Aging clocks reveal a rejuvenation event during early embryogenesis. **(A) - (D)**, The same analysis as shown in fig. S3, but separated by sample type (instead of embryonic day). Epigenetic age of gametes and adult samples (heart, brain and liver) are also displayed.

**Figure S5.**
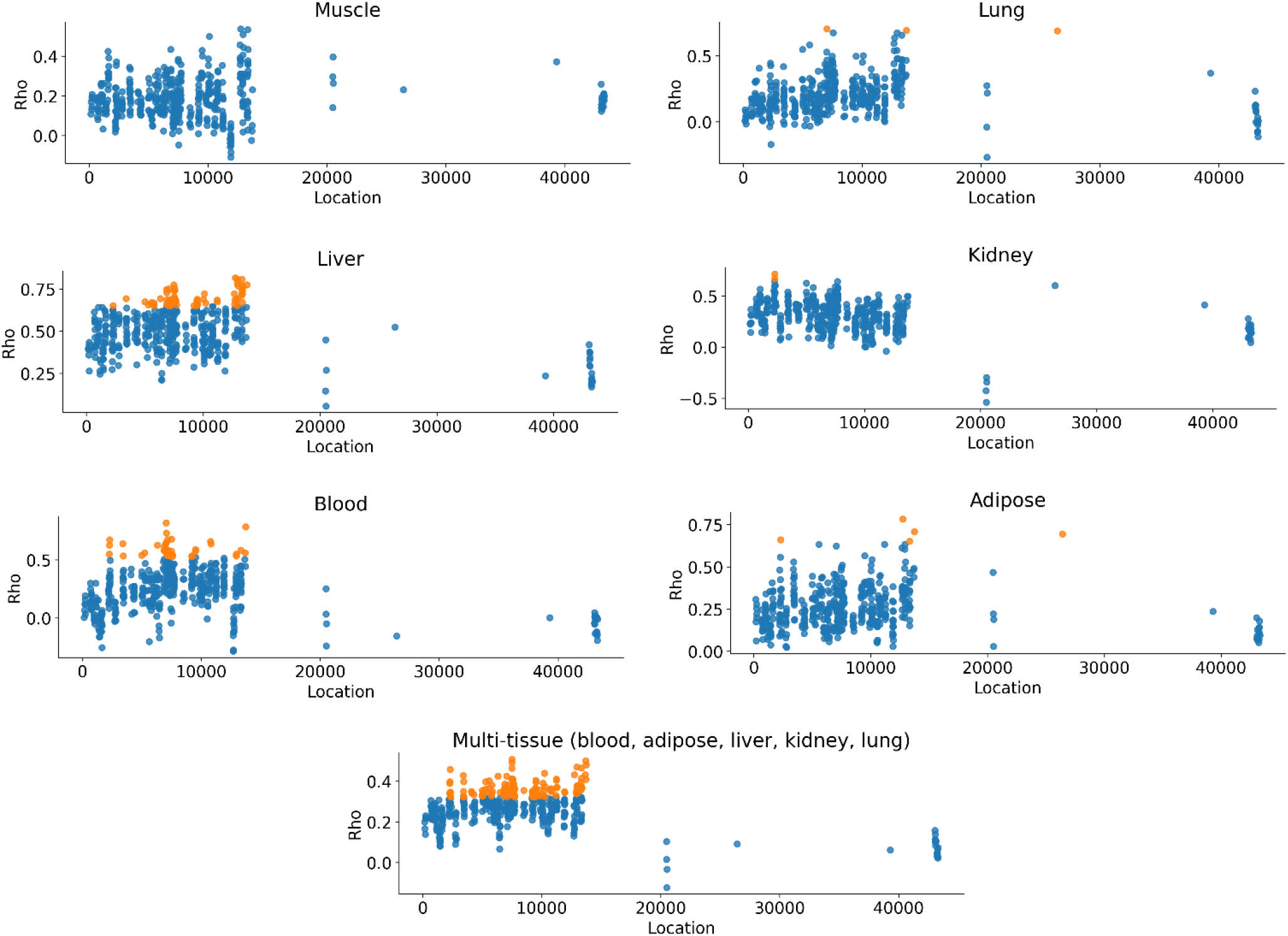
Ribosomal DNA methylation is associated with age in different tissues of C57BL/6J mice. Spearman’s rank correlation coefficients with age (Rho) for CpGs sites of the rDNA sequence. Significant correlations after Bonferroni correction (p <= 0.05 / n) are indicated by orange color.

**Figure S6.**
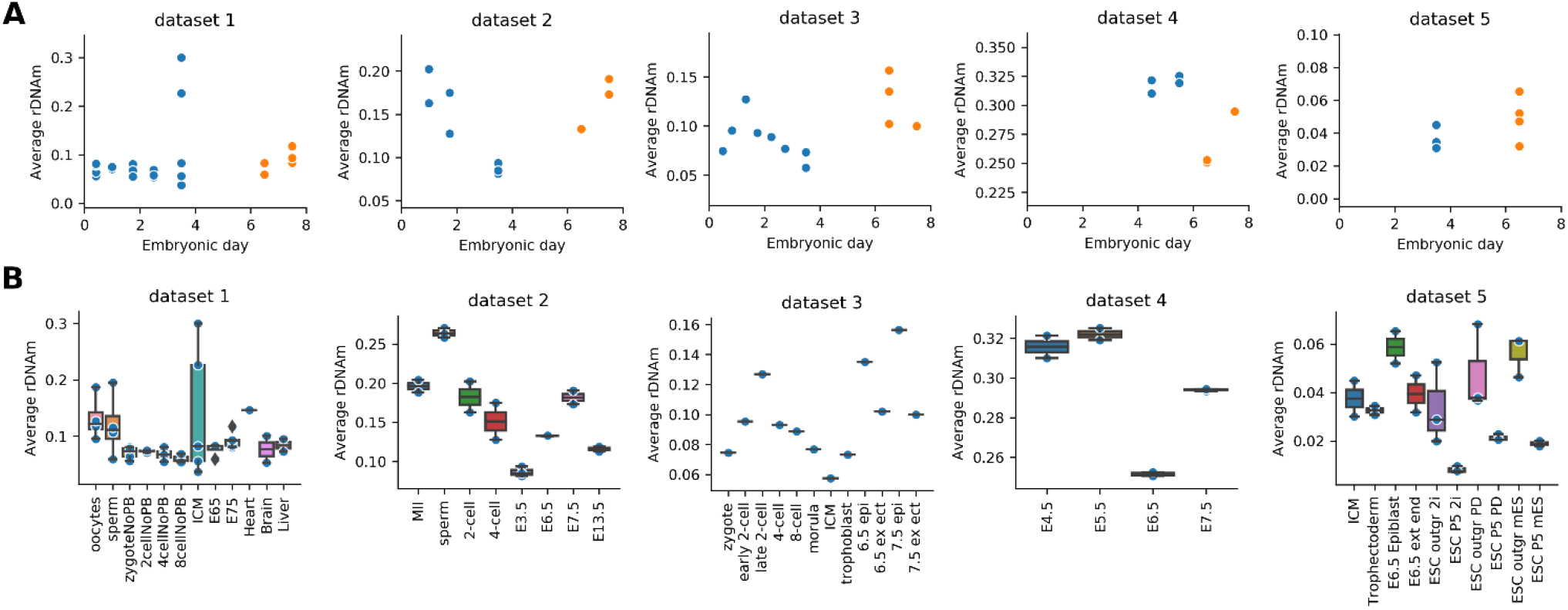
Average methylation level of CpG sites of mouse ribosomal DNA measured in different datasets. **(A)** Samples are separated by embryonic day. **(B)** Samples are separated by type.

